# The SARS-CoV-2 Spike Protein Activates the Epidermal Growth Factor Receptor-Mediated Signaling

**DOI:** 10.1101/2022.05.10.491351

**Authors:** Abdulrasheed Palakkott, Aysha Alneyadi, Khalid Muhammad, Ali H. Eid, Khaled Amiri, Mohammed Akli Ayoub, Rabah Iratni

**Author notes:** The authors contributed equally to the work (Abdulrasheed Palakkott, Aysha Alneyadi). Correspondence: Mohammed Akli Ayoub, Rabah Iratni.

## Abstract

**Objectives:** The coronavirus disease-19 (COVID-19) pandemic is caused by the novel severe acute respiratory syndrome coronavirus 2 (SARS-CoV-2). At the molecular and cellular levels, the SARS-Cov-2 uses its envelope glycoprotein, the spike S protein, to infect the target cells in the lungs via binding with their transmembrane receptor, the angiotensin-converting enzyme 2 (ACE2). Here, we wanted to invesitgate if other molecular targets and pathways may be used by SARS-Cov-2.

**Methods:** We investigated the possibility for the spike 1 S protein and its receptor-binding domain (RBD) to target the epidermal growth factor receptor (EGFR) and its downstream signaling pathway *in vitro* using the lung cancer cell line (A549 cells). Protein expression and phosphorylation was examined upon cell treatment with the recombinant full spike 1 S protein or RBD.

**Results:** We demonstrate for the first time the activation of EGFR by the Spike 1 protein associated with the phosphorylation of the canonical ERK1/2 and AKT kinases and an increase of survivin expression controlling the survival pathway.

**Conclusions:** Our study suggests the putative implication of EGFR and its related signaling pathways in SARS-CoV-2 infectivity and Covid-19 pathology. This may open new perspectives in the treatment of Covid-19 patients by targeting EGFR.

## Introduction

The new COVID-19 disease was identified for the first time in December 2019 in the province of Wuhan in China and it is caused by a new member of coronavirus family called SARS-CoV-2 (Wang et al., 2020; Zhu et al., 2020; Hu et al., 2021). According to the latest statistics, over 111 million COVID-19 cases were reported causing around 2.5 million deaths worldwide (https://www.who.int/emergencies/diseases/novel-coronavirus-2019).

The molecular and cellular basis of the SARS-CoV-2 infection implies the renin-angiotensin system (RAS) and more importantly its angiotensin-converting enzyme 2 (ACE2) (Li et al., 2003; Tolouian et al., 2020; Hoffmann et al., 2020; Yan et al., 2020; Tai et al., 2020; Sriram et al., 2020; Hu et al., 2021). ACE2 is a host membrane-bound metallopeptidase with the catalytic site oriented extracellularly is mostly expressed in lung, heart, kidney, brain, and gut. In contrast to ACE which converts angiotensin I to the active vasoconstrictor, angiotensin II (AngII), ACE2 breaks down AngII to angiotensin-(1–9 and 1–7), which are potent vasodilators, and considered as a negative regulator of RAS (Lavoie & Sigmund, 2003). The implication of ACE2 in COVID-19 is thought to occur mostly in the very early stages of the viral infection and COVID-19 pathology. Indeed, SARS-CoV-2-S protein, the spike glycoprotein (protein S), on the virion surface has been reported to bind the extracellular domain of ACE2 which is used as a co-receptor for target cell recognition and membrane fusion during the infection process (Li et al., 2003; Tolouian et al., 2020; Hoffmann et al., 2020; Yan et al., 2020; Tai et al., 2020; Sriram et al., 2020a; Hu et al., 2021). ACE2 constitutes the main entry gate for other coronaviruses including the SARS-CoV (Yan et al., 2020). In addition, *in vivo* studies showed a nice correlation between COVID-19 infection and the relative expression of ACE2 (positive) and its activity (negative). Furthermore, the receptor-binding domain (RBD) in SARS-CoV-2 S protein has been identified and shown to bind strongly to human and bat ACE2 receptors (Tai et al., 2020). The purified human recombinant RBD showed a potent competitive action on the binding and, hence, the attachment of SARS-CoV-2 RBD to ACE2-expressing cells and their infection by the pseudo virus (Tai et al., 2020; Hu et al., 2021). Thus, the RBD constitutes the most antigenic entity of the S protein used for the development of vaccines to prevent from SARS-CoV-2 infection. Several spike-protein and RBD-based vaccines are in clinical trial (Krammer, 2020).

The disease severity and mortality of COVID-19 have been reported to be increased in patients suffering from other chronic diseases such as cancer, diabetes, hypertension and cardiovascular problems. This was further evidenced in the patients who have been treated with anti-hypertensive drugs such as ACE inhibitors (ACEIs) and angiotensin receptor blockers (ARBs)(Sriram et al., 2020a; Mourad & Levy, 2020; Guo et al., 2020; Gurwitz, 2020; Oliveros et al., 2020; South et al., 2020; Patel & Verma, 2020; Sommerstein et al., 2020). In parallel, studies have showed that ACEIs and ARBs resulted in an upregulation of ACE2 and favoring the entry and replication of the virus (Sommerstein et al., 2020). By contrast, SARS-CoV-2 by targeting ACE2 on the target cells causes down-regulation and inactivation of the latter. This downregulation generates an imbalance in favor of an over accumulation of AngII, a potent vasoconstrictor. Such effect was shown to increase the oxidative damage which leads to inflammation and pulmonary fibrosis (Mascolo et al., 2020). Moreover, inflammation, cytokine storm, and thrombosis associated with pulmonary injury constitute other important clinical features of COVID-19 pathology. This suggests the implication of other molecular actors such as the protease thrombin via its proteinase-activated receptors (PARs), the purinergic receptors, cytokine receptors, and lipid mediators. Thus, the inhibition of these pathways has been proposed as a promising therapeutic approach to prevent thrombotic and inflammatory processes during COVID-19 pathology (Sriram et al., 2020b; Lee et al., 2021).

Although large body of evidence using *in vitro* studies and *in silico* data strongly support the thesis that ACE2 is necessary for SARS-CoV-2 entry, still, we cannot rule out that additional factors and/or alternative cell surface receptors may be also implicated in SARS-CoV-2 entry (Zamorano & Grandvaux, 2020). Among these possible receptors, the G protein-coupled receptors (GPCRs) and receptor tyrosine kinase (RTKs) constitutes the valid candidates based on their tissue abundance and pivotal roles in human and animal physiology. Indeed, several previous studies reported the hijacking of GPCRs and RTKs and their function by various pathogens during pathogenesis. This includes microbial pathogens such as bacteria and viruses such as the SARS-CoV and the adrenergic receptor and the epidermal growth factor receptor (EGFR) as the targets (Wiedemann et al., 2016; Venkataraman & Frieman, 2017; Mitchell et al., 2019). Thus, we hypothesize that during SARS-CoV-2 infectivity, the virus may also use EGFR expressed on the epithelial lung cells as the receptor/co-receptor target for its entry. Here, we have examined the effect of the SARS-COV-2 full-length and DRB Spike 1 protein on the activation of EGFR and its related downstream signaling pathways consisting of AKT and ERK1/2 phosphorylation in the lung cancer cells (A549).

## Materials and Methods

### Cell culture, chemicals and antibodies

The lung cancer cells (A549) used in this study were obtained from Cell Line Service (CLS)-GmbH and were maintained in RPMI (Cat. # 00506 Gibco, Life Technologies, Rockville, UK) complemented with 10% fetal bovine serum (FBS) (Cat. # 02187 Gibco, Life Technologies, Rockville, UK) and 100 U/ml penicillin streptomycin glutamine (Cat. # 01574 Gibco, Life Technologies, Rockville, UK). AG1478 (Cat. # 141438) was purchased from Abcam, Cambridge, UK. Antibodies against phospho-EGFR (Cat. # 4407), EGFR (Cat. # 4267), phospho-ERK1/2 (Cat. # 9106), ERK1/2 (Cat. # 4695), phospho-AKT (Cat. # 9271), AKT (Cat. # 9272), Survivin (Cat. # 2803), anti-mouse (Cat. # 7076), and anti-rabbit (Cat. # 7074) were purchased from Cell Signaling, Technology, Danvers, Massachusetts USA. SARS-COV-2 spike protein 1 (Cat. # DAGC091) was purchased from Creative Diagnostics, Shirley, NY, USA. SARS-COV-2 spike RBD protein (Cat. # 40592-V08B) was purchased from Sino Biological, Beijing China. Human EGF Recombinant Protein (Cat. # PHG0313) was purchased from Gibco, Life Technologies, Rockville, UK.

### Whole Cell extract and Western Blotting analysis

Cells (0.5 x 10^6^) were seeded per well in 60 mm culture dish and cultured for 24h before treatment. After treatment with or without EGF, spike 1 protein or Spike RBD protein, cells were washed twice with ice-cold PBS, scraped, pelleted, and lysed in RIPA buffer (Pierce) supplemented with protease inhibitor cocktail (Roche) and phosphatase inhibitor (Roche). After incubation for 30 min on ice, cell lysates were centrifuged at 14,000 rpm for 20 min at 4°C. Protein concentration of lysates was determined by BCA protein assay kit (Thermo Scientific) and the lysates were adjusted with lysis buffer. Aliquots of 25 μg of total cell lysate were resolved onto 6-15% SDS-PAGE. Proteins were transferred to nitrocellulose membranes (Thermo Scientific) and blocked for 1 h at room temperature with 5% non-fat dry milk in TBST (TBS and 0.05% Tween 20). Incubation with specific primary antibodies was performed in blocking buffer overnight at 4°C and this was true for all the primary protein antibodies used in this study. Notice that all the lysates were freshly loaded when different phospho-proteins were analyzed. However, the same membrane of the phospho-protein was stripped to examine the corresponding total protein loaded except for pERK1/2 where the proteins were always freshly loaded due to stronger anti-pERK1/2 binding. Moreover, a β-actin blot was also used in parallel to double check that similar amount of proteins were loaded in every gel’s lane. Horseradish peroxidase-conjugated anti-IgG was used as secondary antibody. Immunoreactive bands were detected by ECL chemiluminescent substrate (Thermo-Scientific) and chemiluminescence was detected using the LiCOR C-DiGit blot scanner and Image Studio Light Software.

## Results

### Spike and RBD activate EGFR, AKT, and ERK1/2 in A549 cells

First, we examined the effect of Spike 1 and RBD on the phosphorylation status of EGFR and its related kinases, AKT and ERK1/2, in lung cancer cells (A549) (**Figure 1**). The treatment of cells for 5 minutes with Spike (2.5 μg/ml) promoted a strong phosphorylation of EGFR, AKT, and ERK1/2, which was either similar (EGFR and ERK1/2) or even higher (AKT) to that promoted by stimulation with the maximal dose of EGF (5 μg/ml) (**Figure 1**). On the other hand, RBD (5 μg/ml) under a similar condition showed almost no effect on EGFR phosphorylation while it significantly induced AKT and ERK1/2 phosphorylation (**Figure 1**). This result suggests the specific targeting of EGFR by the full spike 1 protein of SARS-COV-2 but not by its RBD. Also, our data suggest that RBD-mediated AKT and ERK1/2 activation is independent on EGFR activation and that other molecular targets and/or intracellular mechanisms may be involved.

**Figure 1.**
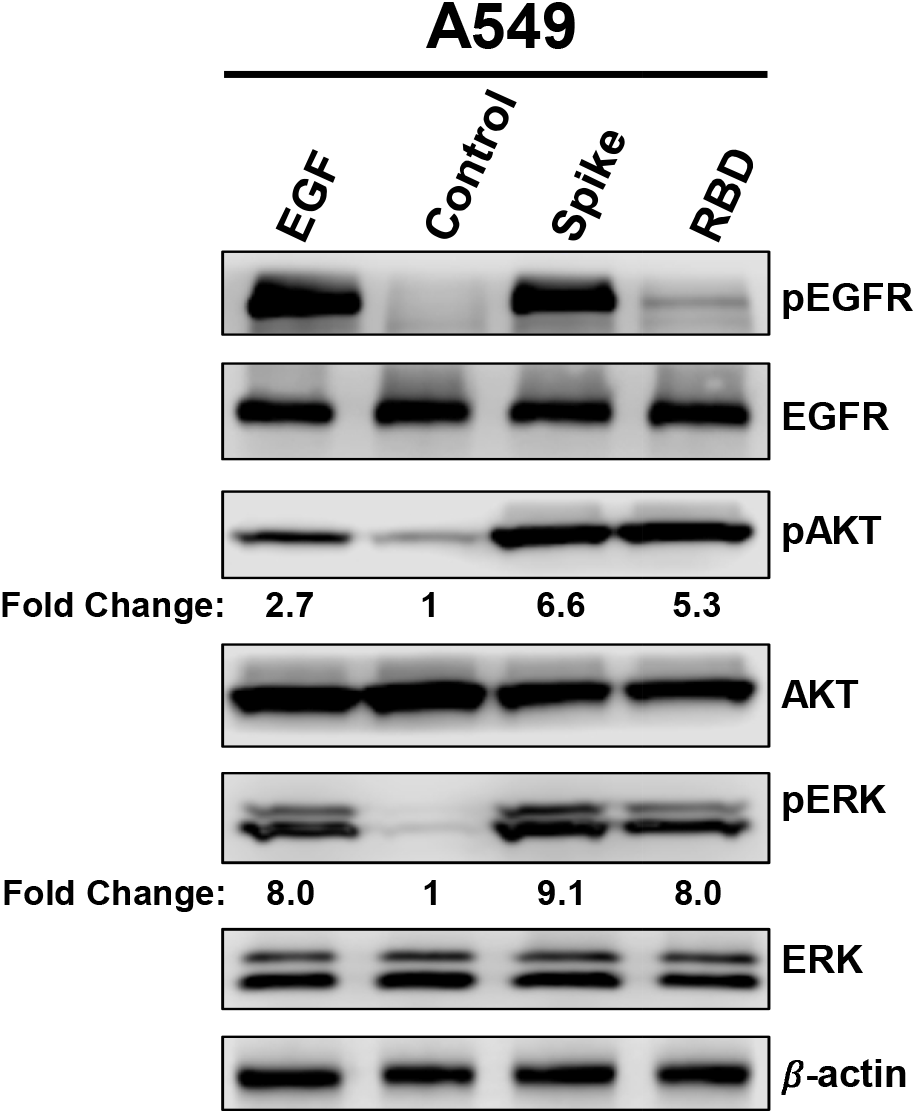
Effect of SARS-COV-2 Spike 1 protein on EGFR, AKT and ERK1/2 activation. A549 cells were treated or not with EGF (5 μg/mL) for 15 min or with full-length Spike 1 (2.5 μg/mL) or with RBD (5 μg/mL) protein for 5 min then whole-cell extracts were subjected to western blot analysis for pEGFR, pAKT, pERK1/2 and their respective total levels.

We next performed time-course analysis with both Spike 1 and RBD on the phosphorylation status of EGFR and ERK1/2. For this, A549 cells were treated with Spike 1 (2.5 μg/ml) (**Figure 2A**) or RBD (5 μg/ml) (**Figure 2B**) for 5, 15, 30, and 60 minutes. Stimulation with EGF (5 μg/ml) as a positive control was carried out for 15 minutes. As shown in **Figure 2A**, Spike 1 induced a rapid and a transient phosphorylation of EGFR and ERK1/2 with a pic at 5 minutes of stimulation and sharp decrease after 15 minutes. Spike 1-mediated AKT activation, on the other hand, was also observed upon 5 minutes of stimulation but this was more sustained in time (**Figure 2A**). For RBD, there was no clear EGFR phosphorylation induced regardless of the stimulation time (**Figure 2B**). This is consistent with our observation in **Figure 1B**. However, RBD promoted rapid and strong activation of both ERK1/2 and AKT that remained persistent even after 60 minutes of stimulation (**Figure 2B**).

**Figure 2.**
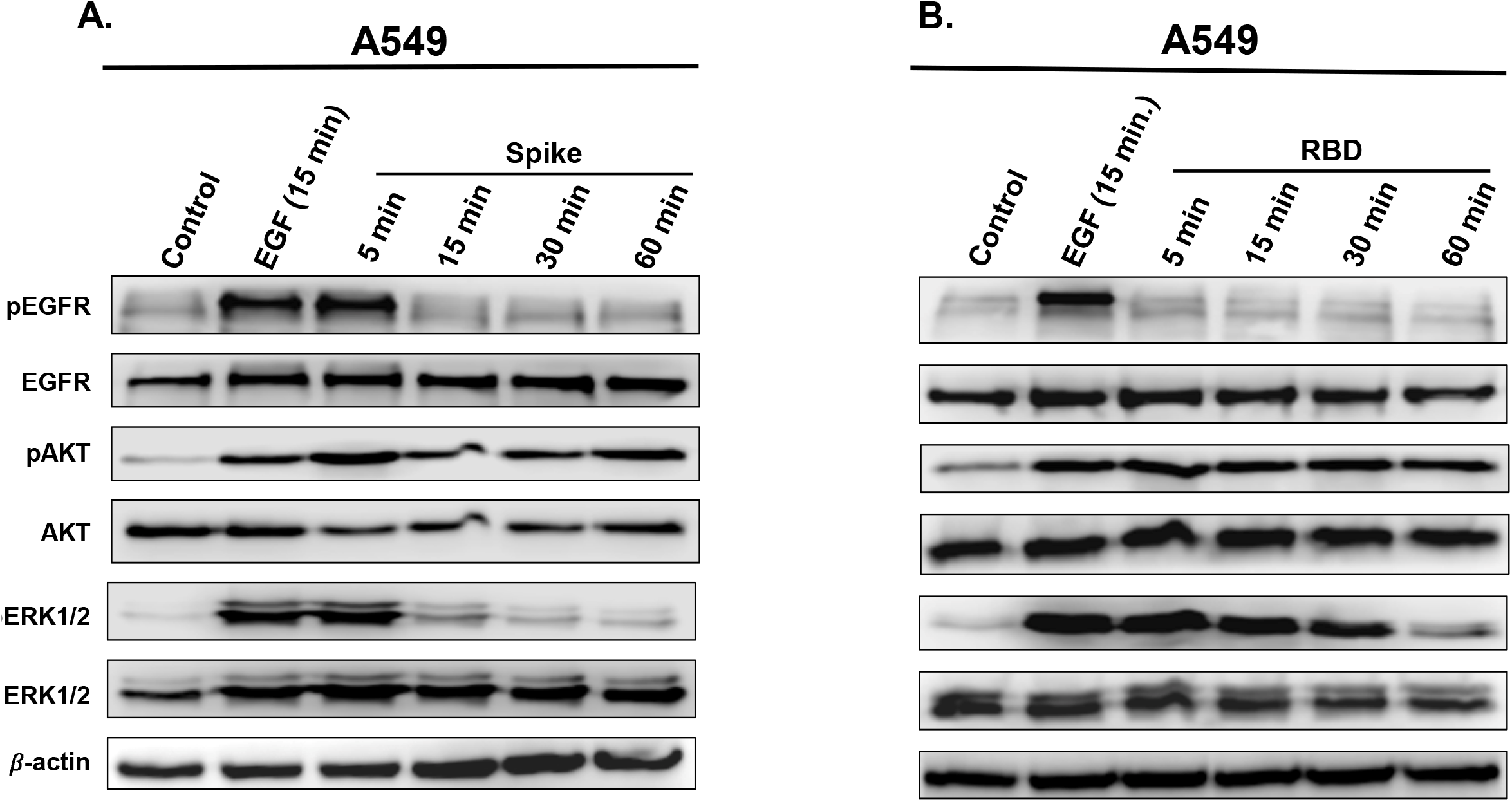
Time-course accumulation of pEGFR and pERK1/2 in A549 cells treated with full length Spike 1 protein and its RBD. Cells were treated or not with 2.5□μg/mL of full length Spike 1 protein (**A**) or 5□μg/mL of RBD (**B**) at different times as indicated (5, 15, 30 and 60 min) and the protein levels of pEGFR and pERK1/2 and their respective total levels were determined by western blot. The treatment with EGF (5 μg/mL) was carried out for 15 min.

Together, these data further confirm the specific activation of EGFR by Spike 1, but not RBD, but they also demonstrate the implication of additional and/or different mechanisms in AKT and ERK1/2 activation whether cells were activated by Spike 1 or RBD.

### Activation of AKT by Spike 1 and RBD is EGFR-dependent

Next, we decided to further explore the targeting of EGFR and its related downstream AKT survival pathway by Spike 1 and RBD by using the selective EGFR antagonist, AG1478. As shown in **Figure 3A**, the pre-treatment of A549 cells with AG1478 (10 μM) fully blocked EGFR phosphorylation induced by stimulation with EGF (5 μg/ml) indicating the specificity of the response. We found that, AG1478 completed abolished AKT phosphorylation induced by EGF, Spike 1, or RBD (**Figure 3B**). This finding strongly suggests that Spike 1- and RBD-mediated AKT phosphorylation is dependent on EGFR activation.

**Figure 3.**
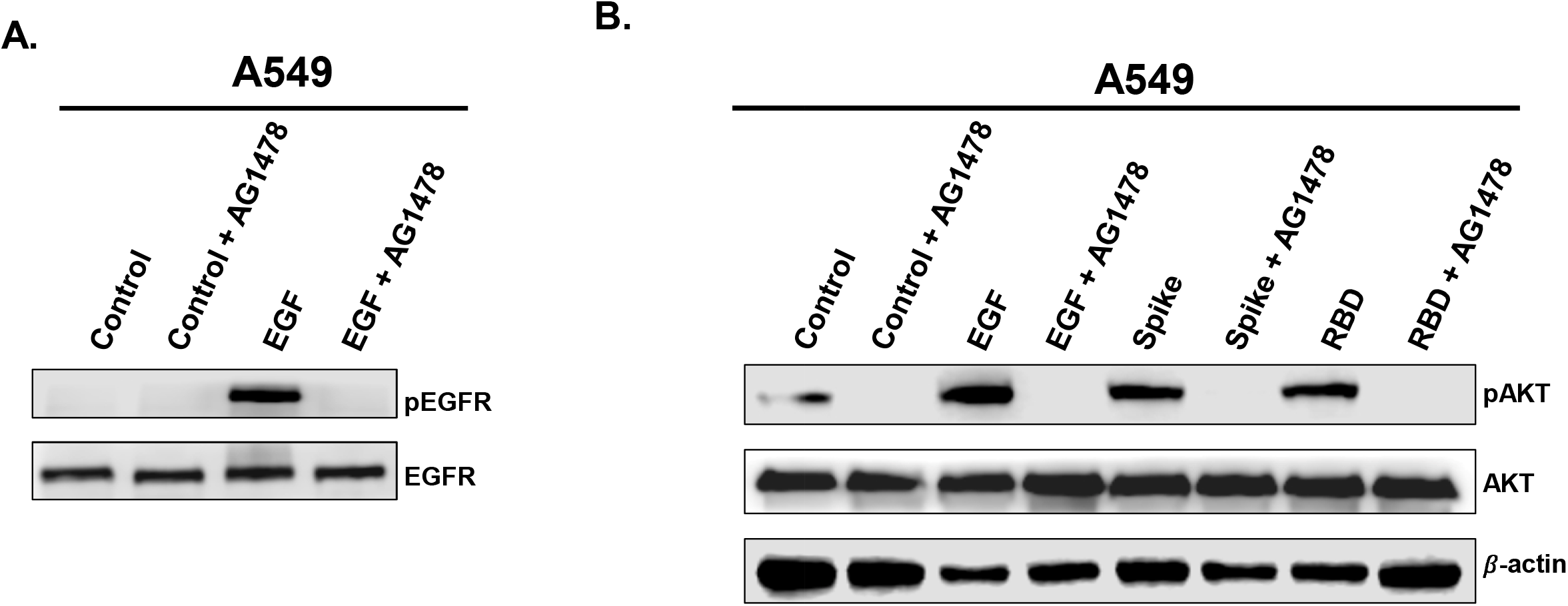
Blockade of EGFR inhibits SARS-COV-2 spike 1-mediated activation of AKT in A549 cells. A549 were first pre-treated with AG1478 (10 μM) for 15 min prior to treatment with EGF (**A** and **B**), full-length spike 1 protein (**B**), or RBD (**B**) as described in **Figure 1**. Whole cell extracts were resolved on SDS-PAGE and protein levels were determined by western blot.

### Effects of Spike 1 and RBD on the cell survival marker, survivin

Since EGFR, AKT, and ERK1/2 pathways are known for their role in cell proliferation and cell survival, we wanted to link our data on the phosphorylation status of these proteins especially EGFR and AKT with the induction of the anti-apoptotic protein, survivin, which belongs to the inhibitor of apoptosis (IAP) family and considered as the key marker for the activation of the survival pathway in cells. Moreover, the activation of survivin was shown to be induced by both AKT and ERK1/2 signaling in cells. For this, we wanted to test whether Spike- and RBD-mediated activation of ERK1/2 and AKT was associated with the induction of surviving expression. Toward this, A549 cells were treated with Spike 1 (2.5 μg/ml) (**Figure 4A**) or RBD (5 μg/ml) (**Figure 4B**) for 5, 15, 30, and 60 minutes using the stimulation 15 minutes with EGF (5 μg/ml) as a positive control and the level of surviving was determined by Western Blot. We found that both Spike 1 (**Figure 4A**) and RBD (**Figure 4B**) induced a marked increase of survivin expression in a time-dependent manner with a maximal response at 30 and 60 minutes. These observations demonstrate that Spike 1- and RBD-promoted EGFR/AKT pathway in A549 cells is associated with the activation of survivin that may promote the survival of the SARS-Cov-2-infected lung cancer cells.

**Figure 4.**
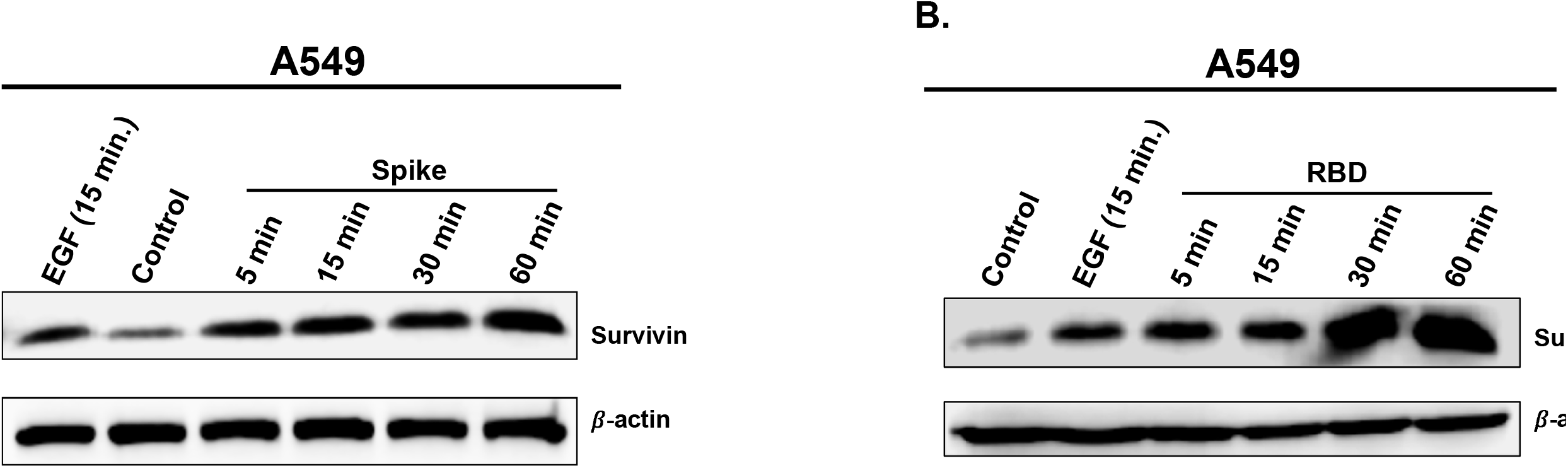
SARS-COV-2 full length spike 1 protein and its RBD promote survivin expression in A549 cells. A549 cells were treated or not with EGF (5□μg/mL) (**A** and **B**), full-length spike 1 protein (2.5□μg/mL)(**A**), or RBD (5□μg/mL)(**B**) at different times (5, 15, 30 and 60 min) as indicated and the protein level of survivin was determined by western blot.

## Discussion

In this study, we report for the first time the possible targeting of EGFR and its related downstream signaling pathways by SARS-Cov-2 Spike 1 protein and its RBD in lung cancer cells (A549). We demonstrated that Spike 1-induced AKT activation occurred in EGFR-dependent manner since it was drastically blocked by AG1478. On the other hand, we found that Spike 1 and RBD also elicited the activation of the survival pathway in A549 cells. Indeed, both Spike 1 and RBD induced the expression and the activation of the anti-apoptotic protein, survivin, which belongs to the inhibitor of apoptosis (IAP) family and considered as the key marker for the activation of the survival pathway in cancer cells. Such a response was very consistent with the phosphorylation of AKT in these cancer cells. This may constitute a solid molecular and cellular rationale to explain the increased risk of infectivity by SARS-Cov-2 and its severity in cancer patients as recently reported by several groups (Albiges et al., 2020; Dai et al., 2020).

Regarding the implication of our findings in the pathophysiology of COVID-19, recent studies showed that cancer patients were more vulnerable to the SAR-COV-2 infection. Although COVID-19 was reported to have low death rate ~2% in the general population, patients with cancer and COVID-19, have at least 3-fold increase in the death rate. Dai et al., 2020 showed that patients with lung cancer, gastrointestinal cancer or breast cancer had the highest frequency of critical symptoms including highest death rates. Patients with lung cancer and gastrointestinal cancer had a death rate of 18.18% and 7.69%, respectively (Dai et al., 2020). Interestingly, they showed that cancer patients that received targeted therapy that includes the EGFR-tyrosine kinase inhibitors showed the lowest death rate compared to cancer patients who received immunotherapy, chemotherapy or surgery (Albiges et al., 2020). Another study carried out in Gustave Roussy Cancer Centre (France) by Albiges et al., 2020 showed that 27% of the cancer patients with COVID-19 developed clinical worsening and 17.4% died (Albiges et al., 2020). Here we showed that SAR-COV-2 activates the EGFR and its downstream signaling pathways controlling cell survival and proliferation. We also found that the inhibition of EGFR abolished the SARS-COV-2 activation of AKT.

At the molecular level, our *in vitro* data provide, for the first time, the evidence that SARS-Cov-2 Spike 1 protein activating EGFR and its downstream signaling pathways, AKT and ERK1/2. This is very consistent with the well-established concept of the hijacking of cell surface receptors and their activity/signaling by pathogens including viruses (Sodhi et al., 2004; Zhang et al., 2016;; Ranadheera et al., 2018; Mitchell et al., 2019) and bacteria (Coureuil et al., 2010; Wiedemann et al., 2016). This implies that pathogens use GPCRs and RTKs at the cell surface of the target cells during the infection process leading to their entry in the target cells. Interestingly, previous studies also showed the role of EGFR and its downstream signalling pathways in viruses/bacteria pathogenicity being consistent with our findings on SARS-Cov-2 spike protein (Mitchell et al., 2019; Wiedemann et al., 2016). Indeed, EGFR was shown to be important during influenza infection (Mitchell et al., 2019). In addition, similarly to our data, the Salmonella Rck membrane protein has been reported to bind and to activate EGFR and its mediated signalling resulting in receptor/bacteria co-internalization and cell infection (Wiedemann et al., 2016). This also occurs with GPCRs since both viruses and bacteria were demonstrated to bind and activate GPCRs resulting in the co-internalization of viruses (Sodhi et al., 2004) or bacteria (Coureuil et al., 2010; Wiedemann et al., 2016) with the target receptors. As stated above, we showed that the Spike 1-induced AKT activation occurred in EGFR-dependent manner in A549 cells since it was blocked by EGFR blockade (AG1478). This supports the conclusion that the activation by Spike 1 of the AKT/survival axis depends on EGFR activation (**Figure 5**). Moreover, the differential effects of Spike 1 and RBD suggests different mode of activation and/or molecular pathways involved. One possible explanation is that Spike 1 may directly target EGFR while RBD uses other targets at the cancer cell surface including ACE2 resulting in AKT and ERK1/2 phosphorylation independently on EGFR (**Figure 5**). Overall, our data are consistent with direct targeting of EGFR/AKT pathway, but this does not rule out the alternative pathway consisting of the implication of the canonical ACE2 pathway via transactivation of EGFR at the cell surface or intracellular crosstalk between their intracellular pathways (**Figure 5**). Of course, further studies are required to demonstrate whether Spike 1 protein directly binds to EGFR or not and what would be the implication of the canonical ACE2 pathway. Even though, we did not investigate all the aspects of EGFR-dependent pathways and SARS-Cov-2 infectivity, we believe that our data will pave the way towards further investigation the exact role of EGFR in SARS-Cov-2 infection and pathogenicity.

**Figure 5.**
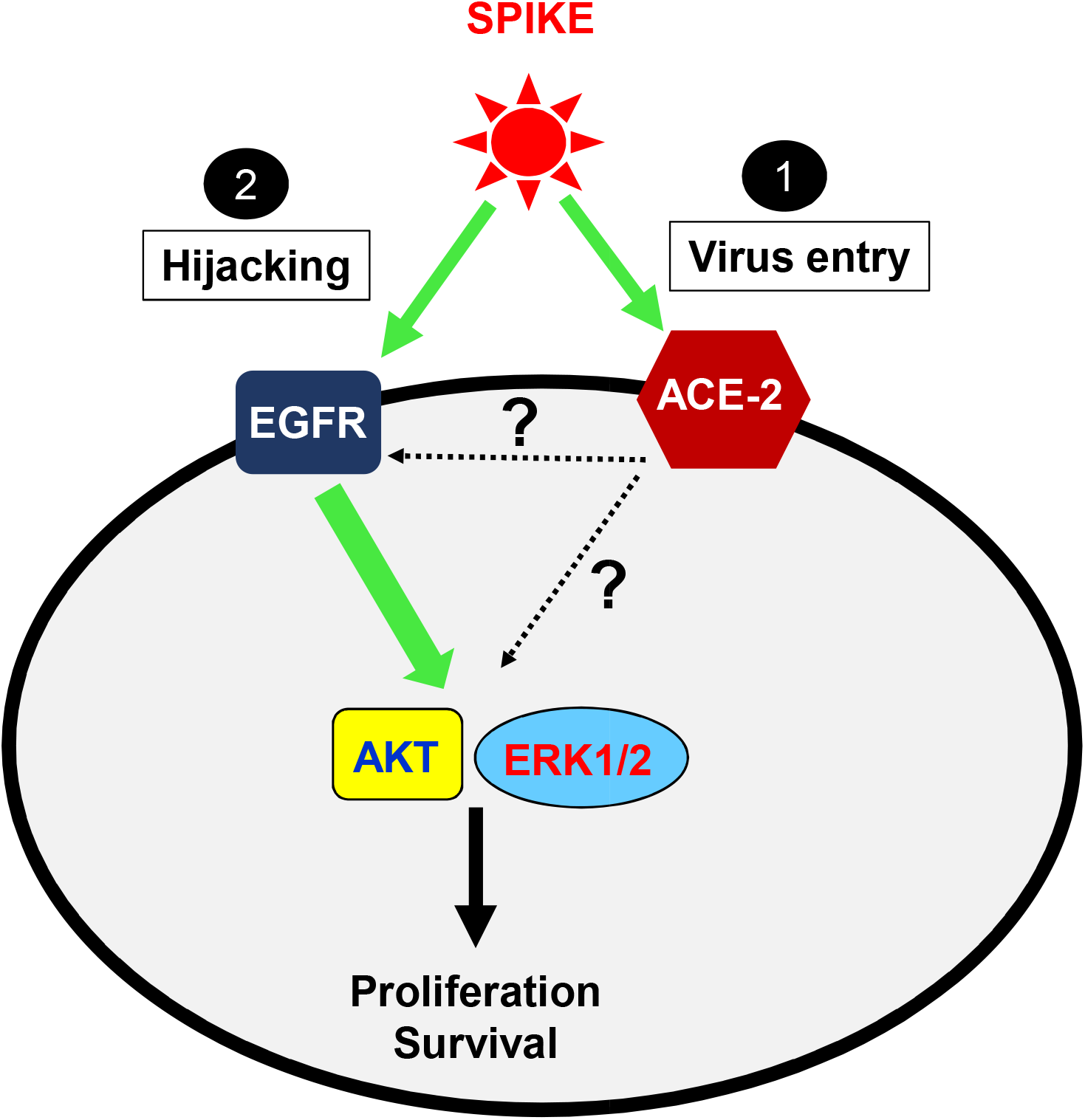
Speculative model for the targeting of EGFR and its downstream signaling by SARS-COV-2 spike 1 protein in A549 cells.

## Author Contributions

AP, HA, MK performed the Western blot experiments; KA and AEH helped analyzing the data; RI and MAA designed the project, analyzed the data and wrote the manuscript. All authors reviewed the manuscript.

## Ethical approval statement

The authors declare no ethical approval was needed for this study.

